# KOMB: Graph-Based Characterization of Genome Dynamics in Microbial Communities

**DOI:** 10.1101/2020.05.21.109587

**Authors:** Advait Balaji, Nicolae Sapoval, Charlie Seto, R.A. Leo Elworth, Michael G. Nute, Tor Savidge, Santiago Segarra, Todd J. Treangen

## Abstract

Characterizing metagenomic samples via kmer-based, database-dependent taxonomic classification methods has provided crucial insight into underlying host-associated microbiome dynamics. However, novel approaches are needed that are able to track microbial community dynamics within metagenomes to elucidate genome flux in response to perturbations and disease states. Here we describe KOMB, a novel approach for tracking homologous regions within microbiomes. KOMB utilizes K-core graph decomposition on metagenome assembly graphs to identify repetitive and homologous regions to varying degrees of resolution. K-core performs a hierarchical decomposition which partitions the graph into shells containing nodes having degree at least K, called K-shells, yielding *O*(*V + E*) complexity compared to exact betweenness centrality complexity of *O*(*V E*) found in prior related approaches. We show through rigorous validation on simulated, synthetic, and real metagenomic datasets that KOMB accurately recovers and profiles repetitive and homologous genomic regions across organisms in the sample. KOMB can also identify functionally-rich regions in Human Microbiome Project (HMP) datasets, and can be used to analyze longitudinal data and identify pivotal taxa in fecal microbiota transplantation (FMT) samples. In summary, KOMB represents a novel approach to microbiome characterization that can efficiently identify sequences of interest in metagenomes.

## Background

Metagenomes are known hotspots for genomic diversity [1, 2, 3]. Communities in metagenomes consist of individual organisms whose genomes are dynamic because of processes such as gene duplication, gene loss/gain, horizontal gene transfer, and gene rearrangements [4, 5, 6, 7]. These dynamic events are a results of complex interactions that underpin the microbiome [8, 9]. Therefore, characterizing metagenomic samples from diverse environments and sample types is essential to understanding community structures, interactions, and underlying functional information [10, 11, 12, 13, 14]. The main approaches to analyze metagenomes include functional characterization and taxonomic classification pipelines [15, 16, 17]. These approaches, while informative, do not necessarily capture the dynamic gene duplication, gene loss/gain or gene transfer activity found in metagenomic samples over time.

In this study, we present KOMB, a novel algorithm for characterizing a metagenome with a particular emphasis on capturing the structure of how repeated elements appear in the community, both within and across microbes. KOMB uses purely sequence level information and does not use a reference database. KOMB begins with a set of partially assembled sequences (unitigs) from he metagenome which it then partitions into hierarchical “shells”, where higher shells contain repeat regions that have high copy number and are more densely concentrated in a few organisms in the communuity. The result is a profile of a given microbiome driven by how genetic repeats are distributed throughout the community.

### Related Work

While certain metagenomic communities including some human body sites [18, 19, 20] are well studied in different pathological conditions, there exists limited information on a plethora of different microbiomes, hindering their characterization [21, 22, 23, 24]. The difficulty in analyzing these metagenomes can often be attributed to the paucity of curated databases and library of reference sequences [25]. The sheer diversity of organisms in these samples that are yet to be identified and annotated further exacerbates this challenge [26, 1].

In order to deal with high-volume metagenomic data from many sample types that may lack an adequate reference, previous efforts have focused on reference-free approaches to quantify variance and diversity. These fall broadly into two classes. First, some methods rely on De Bruijn graphs, assembly graphs, or scaffold graphs to identify sequence-level variation [27, 28, 29, 30]. An overview of the construction of these various graph types including the contributions of this work are illustrated in Figure 1. These approaches characterize samples by relying on popular graph algorithms like betweenness-centrality to identify repetitive contigs, or finding 2-vertex cuts to extract end points of bubbles, or both. In order to reduce the *O*(*V E*) complexity of betweenness-centrality [31, 32, 33], approximation algorithms have sometimes been substituted. Another recent approach has focused on allowing end-users to efficiently query neighbourhoods of interest in metagenomic-compacted De Bruijn graphs, specifically by an indexing approach that approximates minimum r-dominating sets [34]. Though the approximation schemes make calculation more tractable on large metagenomic datasets, its sample wide accuracy and sensitivity may still be sub-optimal [29].

**Figure 1.**
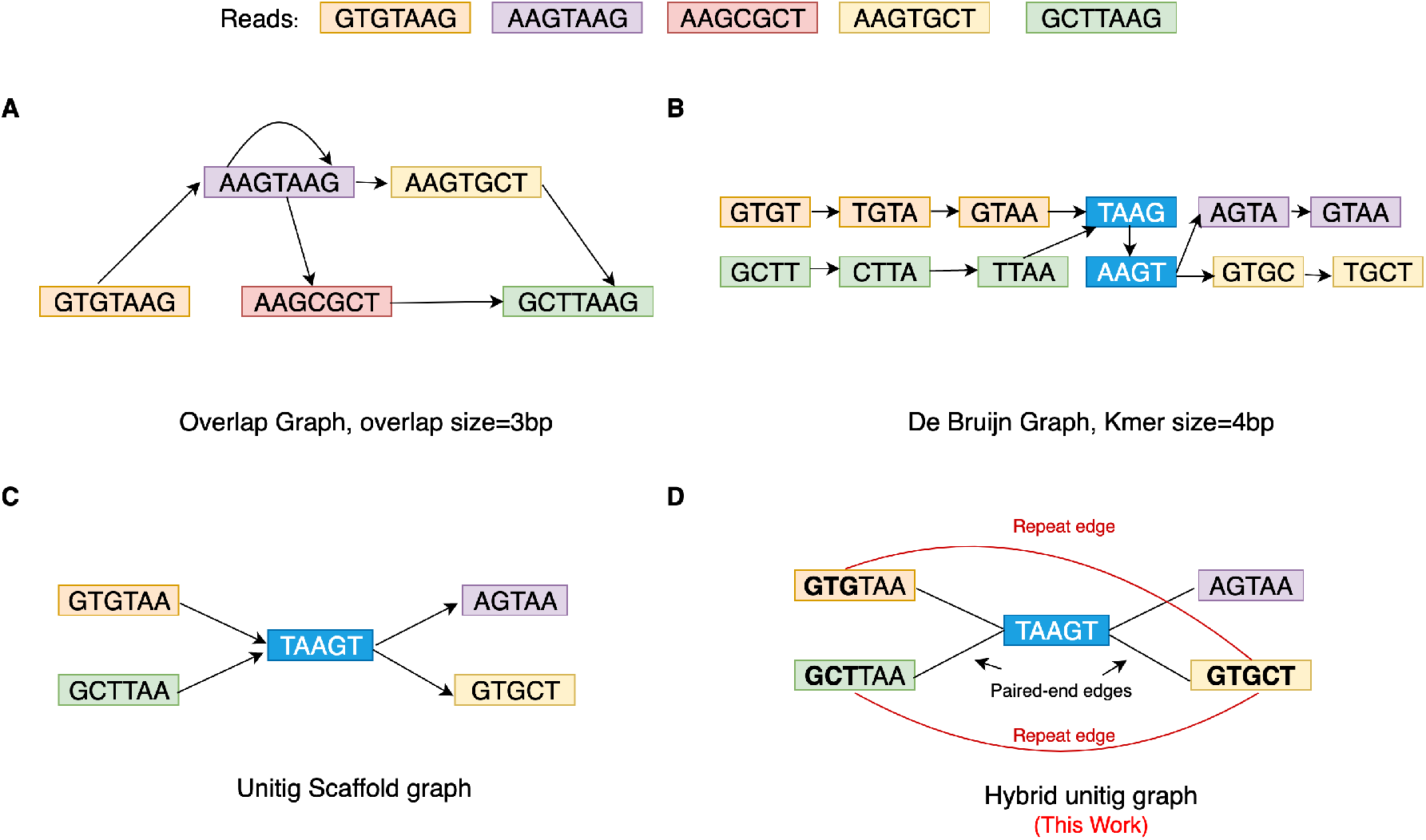
Different graph types for metagenomic analyses and their construction. Graphs construction a set of five reads are shown. A. Overlap graph [Directed]: built directly from read with an overlap size of 3 base pairs(bp). B. De Bruijn graph with kmer size (k) = 4bp [Directed]: joins successive kmers obtained from reads having overlap size of length k-1. The kmers in blue represented repeated kmers. C. Unitig scaffold graph [Directed]: joins unitigs according to their relative positions in a De Bruijn Graph D. Hybrid Unitig Graph [Undirected]: An extension of the Unitig scaffold graph but is also repeat-aware and joins unitigs containing repeats of size k-1 where k is the kmer size used to build the De Bruijn graph. Edge carried forward from the unitig scaffold graph are marked in black and called paired-end edges whereas newly added edges are marked in red and are called repeat edges. 3-mers marked in bold (GTG and GCT) are the repetitive regions connected by the repeat edge.

The second category is k-mer based approaches to quantify diversity and inter-sample distances. These rely on statistical properties of k-mers based on their frequencies [35, 36, 37]. A recent improvement [38] described a generalization of k-mer based method to use Fibonacci Q-matrix in order to efficiently represent every read in the sample as a quadruplet using a sliding-window approach. This matrix was then used for within-sample diversity and inter-sample distance calculations. Though k-mer based methods can efficiently summarize differences up to the sample level, they are not well-suited to identify drivers of genome flux (e.g. duplication or transfer) within a microbial communities.

### KOMB

KOMB is a novel method of characterizing metagenomes that builds off of previous graph-based approaches and incorporates the benefits of k-mer frequency analyses. KOMB relies on the efficient K-core graph decomposition, which has a desirable complexity of *O*(*V + E*). We aim to unify the strengths of graph based and k-mer based approaches to identify both the sequence level features as well as visualize and quantify sample level differences from longitudinal data. Thus, we provide an efficient way to extract micro-level (sequence specific) as well as macro-level (inter-sample distances) insights from short-read metagenomic data. The hybrid unitig graph constructed by KOMB tracks both repeats and rearrangements in metagenomes. To demonstrate KOMB’s usability to profile repeats and capture sequence level features, we apply it to simulated and synthetic data with available ground truths. We also run KOMB on HMP data to illustrate its ability to identify sample specific profiles and functionally rich regions. Finally, we also show KOMB’s ability to capture community disruption events as well as identify markers important to community shifts in longitudinal metagenomic samples on gut microbiome and fecal microbiota transplantation (FMT) samples.

## Methods

### KOMB Algorithm

An overview of the KOMB pipeline is given in Figure 2. The main steps are as follows. First a de Bruijn graph is constructed from reads in the sample subject to some initial filters, and unitigs are identified from this graph. Second, reads are mapped back to unitigs and a graph is constructed on the unitigs by linking them together in two different ways using the read-mapping data (called a “hybrid” graph herein and described in additional detail below). Finally, the hybrid unitig graph is partitioned using the K-core decomposition into an ordered group of subsets (called “shells”), where unitigs in higher shells are have a higher copy number and are densely concentrated whereas those in the earlier shells are more ubiquitous among the organisms in the community. This set of shells along with the unitigs contained in each one is called the KOMB profile, and in what follows we show that it captures a meaningful property of the community.

**Figure 2.**
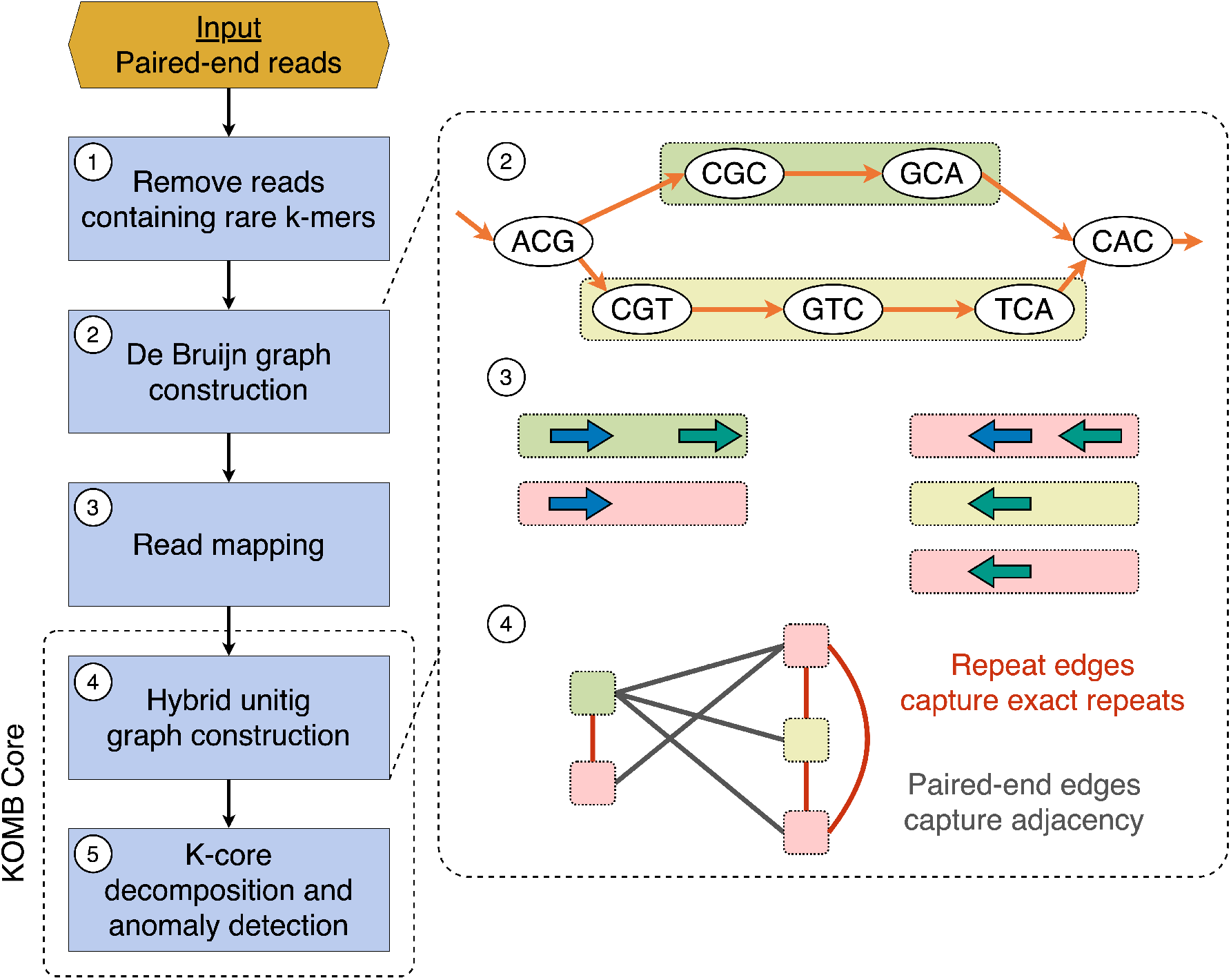
Overview of the KOMB pipeline. 1. As a pre-processing step users can use k-mer filtering t o d iscard l ow-quality e rroneous r eads. 2. K OMB u ses A BySS f or m emory efficient De Bruijn graph construction and unitig generation 3. Paired-end reads are mapped back to the unitigs obtained in 2 in order to connect unitigs. Paired-end reads with just one read mapping are discarded. 4. The hybrid unitig graph is constructed. Edges connecting unitigs mapped by the same read are termed as repeat edges whereas edges between unitigs mapped by paired-end reads are called paired-end edges. The latter are similar to edges in a scaffold graph. 5. The obtained unitig graph is partitioned into K-shells using the K-core decomposition algorithm. Anomalous unitig are marked using the CORE-A anomaly score algorithm.

KOMB incorporates three widely-used bioinformatics tools as part of its workflow. Raw paired-end reads are input to ABySS [39] for efficient De Bruijn graph creation and unitig construction, as well as Bowtie 2 [40] for fast and accurate read mapping. In addition to this, our tool also relies on the igraph C [41] and OpenMP [42] libraries for the K-core implementation and the fast parallel construction of the hybrid unitig graph, respectively. A k-mer based read filtering tool [43] is also available for use as part of the software for optional pre-processing of reads.

### Hybrid unitig graph construction

KOMB constructs a novel hybrid unitig graph to efficiently mine repetitive topologies using K-core graph decomposition. The workflow consists of DBG construction, read mapping, and the KOMB core module as shown in Figure 2. All reads are initially input to the DBG constuctor ABySS to obtain unitigs. A unitig is a maximal consensus sequence usually obtained from traversing a De Bruijn graph. By definition, unitigs terminate at branches caused by repeats and variants and, unlike contigs, are non-overlapping. Subsequently, all of the reads are mapped to unitigs using Bowtie 2. We then construct our hybrid unitig graph with two distinct set of edges. First, for each read we create a set of all unitigs that mapped to that read and connect them. We denote these edges as repeat edges, which capture repeats in unitigs. Second, for a given forward and reverse read pair, we check if each individual read in the pair mapped to different unitigs, which would represent potentially adjacent unitigs in the genome. We call these adjacency edges that attempt to capture any gene loss/gain events between adjacent unitigs. This incorporates paired-end edge information similar to those found in canonical scaffold graphs.

### K-core decomposition

K-core decomposition is a popular graph-theoretical concept used in network science to identify influential nodes in large networks [44, 45, 46]. The K-core of a graph is defined as the maximal induced subgraph where every node has (induced) degree at least *K*. A node belongs to the K-shell if it is contained in the K-core but not in the (*K* + 1)-core. For any given graph, one can iteratively and efficiently decompose it into shells with a complexity proportional to the size of the graph, which is significantly faster than the computation of most exact centrality measures [47]. The shells output as a result of K-core decomposition on the hybrid unitig graph reveal unitigs that are connected either to a similar number of unitigs as a result of their repeat content (via repeat edges) or are adjacent to unitigs with the same properties. At higher shells we observe clique or clique-like behaviours that capture unitigs containing repeats with very high copy number and in some cases appearing very close to each other (e.g., tandem duplications). Both adjacency edges and pseudo-edges are weighted equally in the graph. A more detailed description of K-core decomposition as well as theoretical analysis of the KOMB K-core profile can be found in **Supplementary Figures S1 and S2**.

### Identifying anomalous unitigs

Identification of biologically important unitigs in a given sample is done through ranking the nodes with a CORE-A anomaly score [48]. The CORE-A anomaly score calculates the deviation from mirror pattern (dmp) as given in Equation 1 where *rank*_d_ and *rank*_c_ denote the rank of degree and coreness (shell that a vertex belongs to). This has been shown to reveal nodes of interest in real-world graphs like social and information networks [48]

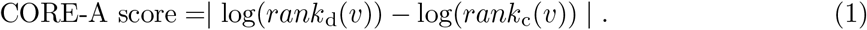

### Datasets

We tested KOMB on four different datasets to illustrate various properties of KOMB and underline different use cases while analyzing metagenomes. The datasets and their use cases are briefly described as follows:

1. **Shakya synthetic metagenome:** A well-characterized synthetic metagenome consisting of 64 organisms (48 bacteria and 16 archea)[49]. This dataset is a simple test case to demonstrate how KOMB operates in practice, how to interpret the results, and how the higher shells reflect the structure of repeated regions in the metagenome.
2. **Multi-site HMP samples:** This dataset contains 50 samples each from four body sites drawn from the Human Microbiome Project (HMP)[50]. These samples are a useful test case for KOMB because they demonstrate a) that the KOMB profile for samples within a given site are broadly similar to one another, and b) that the overall profile for each body site is characteristic and distinct from other body sites in much the same way that the taxonomic profile is. In other words, it suggests that the KOMB profile is both reproducible and is consistent with what might be expected on highly dissimilar communities. This dataset is also used as an example of how the KOMB profile specifically recovers functionally rich sequences.
3. **Longitudinal gut microbiome samples:** This data is also from a previous study [51] and contains samples taken from 6 subjects over two years, including one subject that was exposed to antibiotic and bowel cleanse disruption in that time. This is meant to go one step further by showing that the KOMB profile can capture both subject-specific differences at a common body site and variations in an individual community over time as it is subject to perturbations.
4. **Fecal microbiota transplantation (FMT):** This data has not been previously published and includes samples from two patients undergoing (FMT) from a common donor. Specifically, the samples include both pre- and post-FMT from each patient as well as one sample from the donor. Anomalous unitigs identified in KOMB profiles capture specific taxa that are known to be contributurs to recovery and transition to a disease-free state in Post-FMT samples when compared to both Pre-FMT and Donor samples.

### Running KOMB

The following sections contain detailed descriptions of how KOMB was run on each of the four datasets as well as any steps required for additional analyses discussed in Results below.

### Shakya synthetic metagenome

Reads from the Shakya et al. (2014) study were obtained from NCBI SRA (SRR606249). Reads were filtered using the kmer-filtering tools packaged as part of Stacks [43]. Ground truth for repetitive unitigs was established by using *nucmer* to map the unitigs to the reference genomes with parameters -c 50 -l 50 as the hybrid unitig graph was built on matching 50bp exact matches. KOMB was run with the parameters -k (kmer-size) 51 and −l (read length) 101. Fraction of repeat unitigs were calculated by dividing the number of unitigs marked as repetitive by nucmer to the total number of unitigs in the shell. KOMB repeat density calculated for each shell is given by the formula outlined in Equation 2. We calculate the sum of copy numbers of each repetitive unitig and then divide it by the number of reference genomes these unitigs map to (number between 1-64). This number is then averaged over the number of repetitive unitigs in a shell.

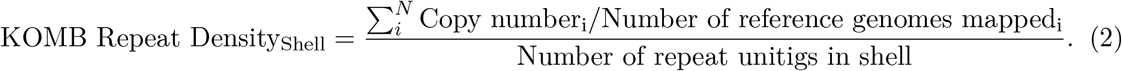

### Multi-site HMP samples

HMP 1 data consisting of 50 samples each from four different body sites (anterior nares, stool, supragingival plaque, and buccal mucosa) was downloaded from the HMP website https://www.hmpdacc.org/HMASM/. Prior to running KOMB, we implemented a homogenizing step where only reads having length equal to the longest read length per sample were kept (mostly 100 bp) and the rest were discarded. KOMB was then run with the parameter -k (kmer-size) 51. Functional characterization of unitigs obtained and marked from the anomaly detection stage is done through SeqScreen [52, 53]. Anomalous unitigs are determined by considering all unitigs whose dmp score (see Equation 1) is above a cutoff score as determined in Equation 3. In this equation, Q_3_ represents third quartlie and I.Q.R is the inter-quartile range which is the difference between the third and first quartiles (Q_3_ - Q_1_). For the analysis, we combined the anomalous unitigs from each individual sample and, separately, we combined the rest of the unitigs from each of the samples to obtain the set of unique GO terms and set of anomalous GO terms for each body site. Anomalous GO terms refers to the GO terms found in unitigs marked as anomalous by KOMB. Unique GO terms refers to a subset of GO terms found only in the anomalous unitigs but not found in other unitigs in a given body site. In other words, anomalous GO terms are a superset of unique GO terms. All GO terms are filtered for bacterial specific GO terms using the https://github.com/AstrobioMike/CoV-IRT-Micro python package. Only GO terms belonging to the *Biological Process* branch were considered for the analysis.

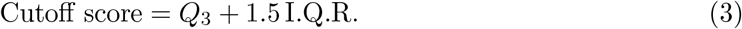

### Longitudinal gut microbiome samples

Reads for the dataset were obtained from the ENA website (ID: ERP009422). The reads were filtered using the kmer filter tool packaged as part of Stacks [43]. The reads were run with the commands -k (kmer-size) 35 and -l (read length) 80.

### Fecal microbiota transplantation (FMT) samples

#### Sample Collection

Two pediatric patients with a recurrent CDI diagnosis received FMT under IRB-approved informed consent (#H-31066) at Baylor College of Medicine. The investigational nature of FMT was highlighted during consenting in accordance with current U.S. Food and Drug Administration (FDA) regulations. CDI diagnosis was based on toxin PCR positivity along with clinical complaints of 3 or more diarrheal stools per day. Patients reported recurrent (return of symptoms within 2 months) or ongoing diarrheal symptoms despite completing at least two courses of CDI-directed antibiotics that included at least one course of metronidazole and vancomycin. Patients received filtered, frozen-thawed fecal preparations from a standardized donor (38-40 y male during donations) via colonoscopy. The donor screening and fecal preparation procedures were approved by the U.S. Food and Drug Administration (IND15743). Fecal samples were collected from patients the day prior to FMT and 8-9 weeks following treatment on a follow-up visit. All samples were frozen and kept at −80°C until simultaneously thawed for bacterial DNA extraction using the PowerSoil DNA isolation kit (MO BIO Laboratories, Carlsbad, California, USA). Shotgun metagenomic sequencing was performed with >200 ng of input DNA as previously described by us [54] and sequence is submitted to NCBI BioProject database: PRJNA743023.

#### Analyses

Reads were mapped to GRCh38p12 using bowtie 2.3.5; with preset options bowtie2 –local; read pairs were extracted from resultant SAM file using samtools 1.9 [55] using flags samtools fastq -f 13; these read pairs were then subjected to KOMB using -k 51 -l 150. Taxonomic analysis of the anomalous unitigs was done by running the unitigs through Kraken2 [56]. Kraken2 was run with the miniKraken2 database v1 (8GB). Unitigs that were successfully classified at genus level or below were considered for the analysis. All unitigs classified at species level were assigned to their corresponding genus. For each sample, anomalous unitigs are obtained by selecting those whose dmp score (Equation 1) is above the cutoff score in Equation 3. For each genus present in anomalous unitigs, we calculate the *Ratio of Ratios* score for each genus as given in Equation 4, where num_ag_ and num_og_ are the number of unitigs classified at genus *g* in the set of anomalous unitigs and other (background) unitigs, respectively. The denominators refer to the sum of all unitigs of all genus present in the set. The total number of unique genus present in both the sets (anomalous and other) are N_a_ and N_o_, respectively. For the analysis, we selected those genera with the ratio of ratios greater than or equal to one (≥ 1) which we term as over-represented genus in the anomalous unitigs.

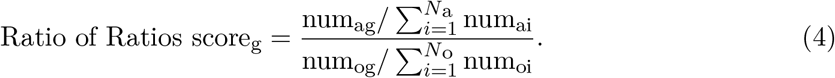

### Calculating the L1 norm between KOMB Profiles

In order to calculate the distance between two KOMB profiles, we use the L1 norm of the difference between their normalized coreness profiles. More precisely, we first divide the size of each shell by the total number of unitigs in each profile. The shorter of the two profiles is then padded with zeros to equalize the number of shells, i.e., we can represent each profile as a vector of the same size. We then compute the distance between the profiles as the L1 norm of the difference between these two vectors.

## Results

### KOMB profile example and interpretation

The Shakya synthetic community was used as a simple example to demonstrate the KOMB profile and provide some evidence to support the assertion that it captures the pattern of repeated regions in the community.

An input to the De Bruijn graph construction is *k*: the exact *k*-mer size used to join reads. The shells in the KOMB profile are labeled incrementally as they are produced in the K-core decomposition. The number of a given shell is approximately the copy number of a family of exact repeats of size *k* − 1 if the unitig is repetitive or the degree of a non-repetitive unitig that is in close proximity to a repetitive unitig with copy number greater than the shell number.

First, Figure 3(A) shows how the full set of unitigs is distributed according to each shell, with a total of 320 K-core shells obtained after decomposition. (For simplicity, we exclude shell 0 which represents isolate unitigs.) Early shells (i.e. 1-4) contain the majority of the unitigs and overall the density declines steeply as the shell number grows, similar to what we might expect in a random graph. However, by contrast with a random graph there are a number of small peaks occurring at higher nodes after the initial drop-off (marked with red triangles). Most of these peaks are followed by regions of empty cores indicating that these peaks mark dense cliques that all share the 50 bp exact match (as 51 was the *k*-mer size used). A similar behaviour was observed in our validation on simulated genomes (Supplementary Figures S3, S4 and S5), where these topological features represented the artificially inserted repeats. These peaks in higher shells are endemic the nature of the KOMB profile, and the number and size of the peaks captured in this figure are a simple summary of the KOMB profile for a given community.

**Figure 3.**
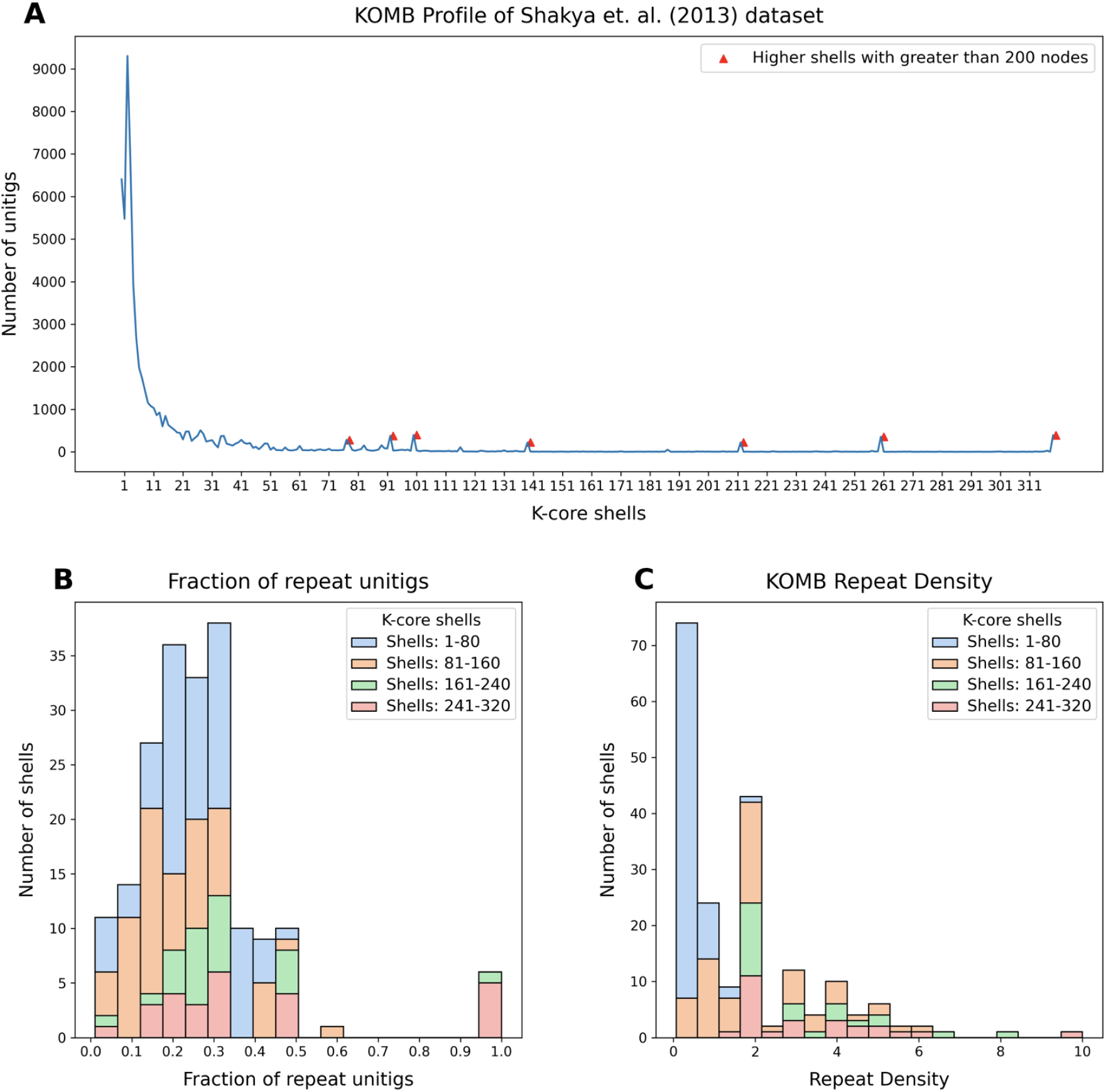
Characterization of a synthetic metagenome sample using KOMB. (A) KOMB profile o f t he S hakya e t a l (2013) d ataset r epresenting t he s hell n umber o n t he x-axis a nd the number of unitigs in the y-axis. Red triangles indicate higher shells with greater than 200 nodes, which represent clique or clique-like regions in the hybrid unitig graph. (B) Histogram representing the fraction of unitigs in each of the shells that are repeats as determined by comparing with the nucmer output. (C) KOMB repeat density is defined a s t he a verage c opy n umber p er n umber of genomes for the repeat unitigs in the shell (see Equation 2). Larger shells have repeats with high copy number but more specific to a single (or group of related) o rganisms. For figures (B) and (C), shell 0 (disconnected nodes) and shells that contained no unitigs are not considered. Shells are split into four different groups (1-80, 81-160, 160-240, 241-320) for visualization.

Beyond simply showing the KOMB profile for this community, it is worth verifying that higher shells do indeed represent regions with more repeats and of higher repeat-number. Here, we have used *nucmer* to quantify the repeat number of each unitig, and the stacked bar charts in Figures 3(B) and 3(C) show how shells compare to one another according to the fraction of unitigs considered a repeat and average repeat density, respectively. The *nucmer* repeat quantification is imperfect and the shells are grouped by quartile, but nonetheless the third and fourth quartiles are skewed to the right in each graph, indicating that indeed the higher shells contain unitigs with a heavier density of repeats. This is a fundamental property of the shells in a KOMB profile. Nucmer analysis of repeat unitigs also revealed that the repeats in the higher shells mapped only to a few organism in the sample but had relatively high copy numbers resulting in a higher density. Combining these observations with those from the KOMB profile, we can infer that the majority of shells containing true repeats are likely to lie beyond shell 161. It is important to note here that the topology of the hybrid unitig graph in addition to repeats the KOMB profile also captures unitigs that are adjacent (as defined by paired-end edges) to repetitive unitigs across copy numbers. Hence, it is expected result that some of shells will not consist a high number of repeats and would instead contain these adjacent unitigs.

### KOMB vis-a-vis beta-diversity and functional annotation

A key test for a novel descriptive profile is whether it is reproducible and whether it shows broad differences where they would intuitively be expected. A key insight about the human microbiome is that the bacterial communities differ substantially by body site, and that communities from the same body site across different individuals are more similar than across body site. We would therefore expect KOMB profiles to follow this same pattern. Figure 4(A) shows the same distribution of unitig density by shell number as in the previous dataset, but here it is presented as a violin plot. Specifically, the plots for all 50 samples from the same site are overlaid to visualize their variability. Each site has its own evident shape, and notably the anterior nares site appears to have the largest range of variability for individual samples. We also analyzed the site-specific profiles for intra-site and inter-site distances which are discussed in **Supplementary Data SD1**.

**Figure 4.**
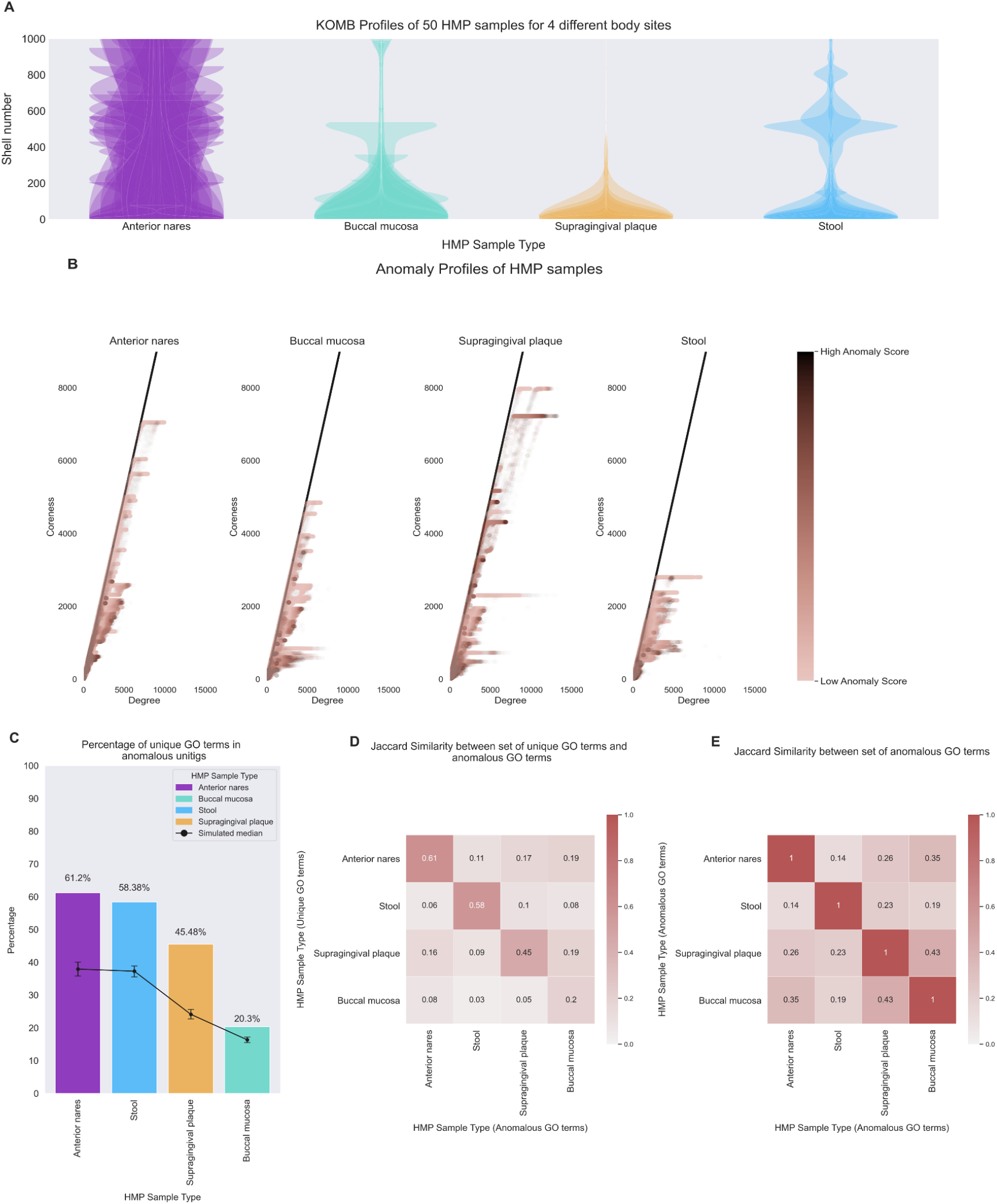
Characterizing community shifts in Human Microbiome Project (HMP) samples. (A) KOMB profiles from 4 different body sites containing 50 samples each obtained from HMP datasets. The y-axis of the violin plots represent shell number (cutoff at 1000 for visualization) and the width represents the number of unitigs in each shell. (B) Anomaly profiles for each body site, x-axis represents the degree of unitigs and y-axis represents the coreness (or shell number) of the unitigs. The gradient on the color bar represents the CORE-A anomaly score with the darker shade representing higher scores within the samples. (C) Bar plot showing the percentage of unique GO terms from the set of unitigs marked as anomalous. Black dots represent median of 100,000 random split simulations of GO terms obtained per body site, the whiskers represent 95*^th^* (top) and 5*^th^* (bottom) percentile indicating significance of the bar plot. (D) Jaccard similarity between the set of unique GO term (y-axis) and the entire set of GO terms from the unitig marked as anomalous for each pair of body sites. (E) Jaccard similarity between the entire set of anomalous GO terms for each pair of body sites.

This dataset also served as a test case for a hypothesis that the KOMB profile could be used to identify highly “important” segments. The K-core decomposition has been useful for this in other contexts, specifically by identifying anomalous nodes in the graph (CITE). Here, we hypothesize that “importance” of a unitig could be represented by functional richness.

We utilize the anomaly detection algorithm as proposed by [48]. Figure 4(B) shows the Coreness vs Degree graph of the unitigs for each body site. The color gradient indicates the CORE-A score with the unitigs having high CORE-A score mainly being high coreness and low degree or low coreness and high degree. Unitigs were separated into those marked as anomalous and those not, then we functionally annotated the unitigs marked as anomalous by assigning GO terms. Then, GO terms occuring only in anomalous unitigs (”unique GO terms”) were expressed as a percentage of all GO terms. For comparison, we conducted simulations in which GO terms were randomly assigned to contigs and ran the same calculation of “unique GO terms”.

Figure 4(C) shows the results: the bar for each body site is the overall % unique, while the black line (and error bars) represent the values obtained by simulation. The actual values are well above the error bars for all body sites, indicating that anomalous unitigs contain a disproportionate share of gene functions that are found *only* in these unitigs. Furthermore, previous studies [57, 58, 59] have described the relative evenness and low diversity of the buccal mucosa community especially in comparison to other oral communities like supragingival plaque which is reflected in our functional analysis of anomalous unitigs.

Further, we analyzed how the unique GO terms in a given body site compare with the GO terms found in anomalous unitigs from other body site. We then calculated the jaccard similarities of these sets. We hypothesized that samples from similar regions (eg. oral) would be more similar functionally than others which we recapitulate in a taxonomy-oblivious manner through KOMB. In Figure 4(D), we see that the jaccard similarities are overall low (< 0.2) indicating that these unique GO terms are generally specific to the microbiome in a given body site. The GO terms in stool were the most dissimilar to those found in anomalous unitigs in other samples (average jaccard similarity=0.05). The similarity scores of unique GO terms in anterior nares and supragingival plaque had greater similarity with the anomalous GO terms buccal mucosa (0.19). Figure 4(E) shows the jaccard similarities of anomalous GO terms between each body site. The oral sites, buccal mucosa and supragingival plaque, had the most similar anomalous GO terms (0.43). Similar to the case with unique GO terms, anterior nares had a higher similarity with buccal mucosa (0.35) than supragingival plaque (0.26). We also observed that anomalous unitigs in stool had the lowest functional similarity to other body sites (average jacaard similarity =0.186). The GO term ID and names can be found in **Supplementary Data SD2**.

### KOMB characterizes community shifts in longitudinal samples Longitudinal gut microbiome samples

To demonstrate KOMB’s ability to derive insights from large scale metagenomic analysis, we considered a temporal gut metagenome study. This study contains git microbiome samples collected from 7 subjects (5 male and 2 female) at different time points spread over two years. Figure 5(A) shows the KOMB profiles of each of the 6 analyzed subjects (one subject was excluded because of missing data point) from the initial three time points (Days 0, 2, 7), each labeled by an alias given in the original study. These violin plots show that the gut samples from the six subjects all have relatively similar KOMB profile distributions, although some idiosyncrasy does appear in subjects Daisy and Bugkiller. To quantify these profiles, The intra-subject and inter-subject sample distances were analyzed and are discussed in **Supplementary Data SD3**.

**Figure 5.**
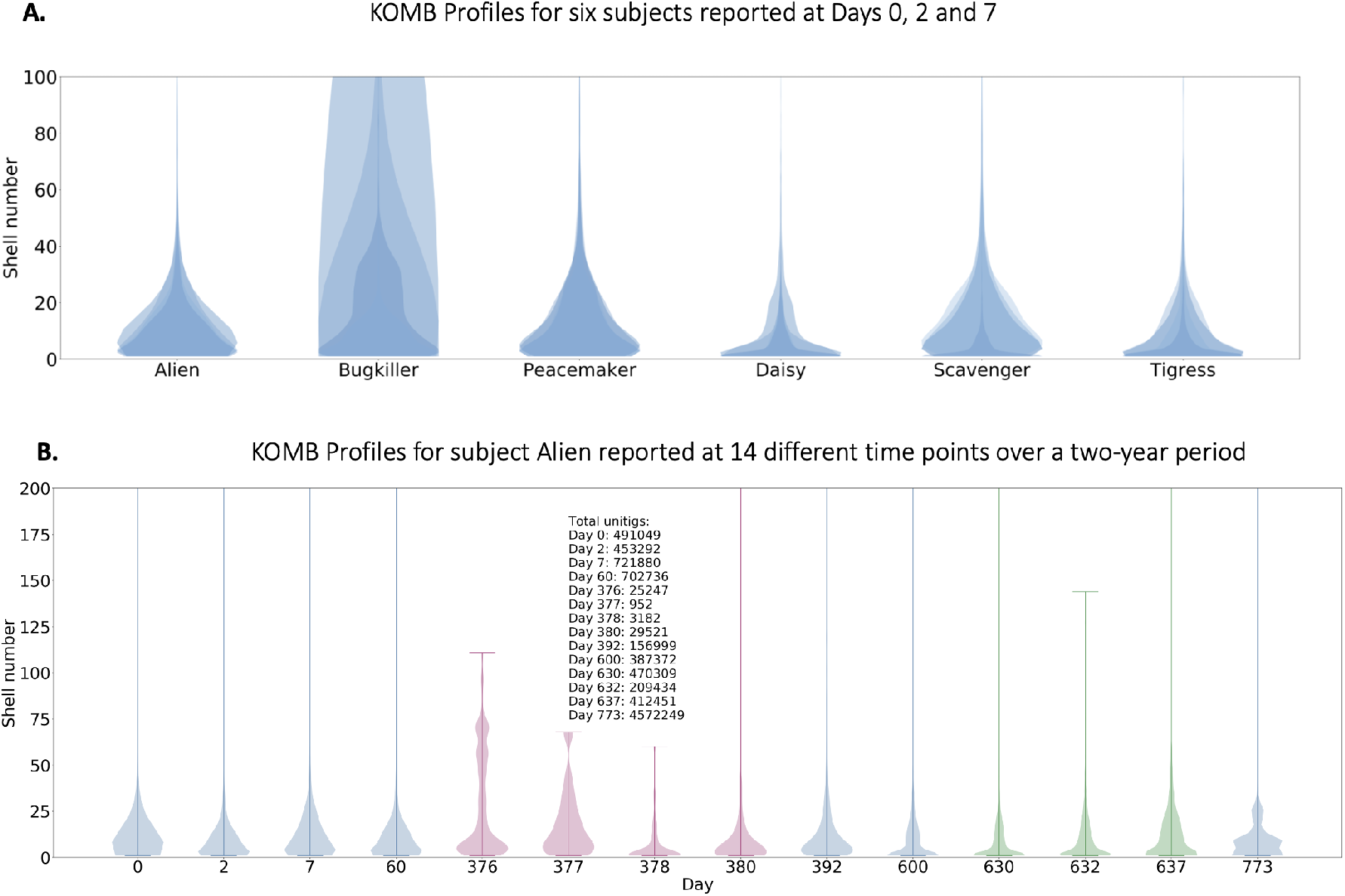
Characterizing community shifts in longitudinal gut microbiome samples. (A) KOMB profiles f rom 6 d ifferent su bjects fr om sa mples co llected Da ys 0, 2 an d 7. Th e y-axis of the violin plots represent shell number (cutoff at 100 for visualization) and the width represents the number of unitigs in each shell. Alien, Bugkiller, Peacemaker, and Scavenger are male subjects while Daisy and Tigress are female subjects. (B) KOMB profile for subject Alien over the course of the 14 different time p oints in the s tudy. The y-axis (cutoff at 200 for visualization) represent shell number and x-axis represents the day of sample collection. Days 376, 377, 378, and 380 represent profiles during which the subject was exposed to a ntibiotics, causing compression in the total shell count as well as a significant change in the unitig distribution of the initial s hells. Days 630 and 632 indicate time points when the subject underwent a bowel cleanse procedure with a similar but less prominent effect on unitig count and distribution.

To get a more quantitative understanding of the data and the effects of external disruptions on the gut microbiome we focus our attention on the subject Alien who was the only subject exposed to an antibiotic intervention and bowel cleanse procedure during the course of the study.

Figure 5(B) outlines the entire longitudinal trajectory of Alien’s gut microbiome over the course of 14 time points spread across two years. The KOMB profiles as displayed focus on the first 200 shells at each time point. We observe a significant change of shape in the profile on Days 376, 377, 378, and 380 which coincides with samples taken after antibiotic intake and which correspond to a significant perturbation community composition as reported in the study. This is also mirrored by the unitig counts in the samples, which decreases by an order of magnitude. Importantly, the total number of reads in the samples from each time point are similar and, hence, the change in unitig count is most likely caused by a shift in the community composition. Thus, antibiotic intervention causes not only a reduction in the total number of shells but also alters the unitigs present in the initial shells, though this tends to recover slightly towards the end of the antibiotic cycle on Day 380. The distirbution of unitigs to shells has returned to form twelve days after the last post-antibiotic sample (Day 392), and the raw number of unitigs has returned to earlier levels by Day 600. We observe similar but less drastic shell compression and quick recovery after a bowel cleanse (Days 630, 632) indicating that antibiotics cause a far greater disruption in microbiome community structure, a finding corroborated by the authors in [51] as well as an earlier study [60].

To further quantify the perturbation, we calculated the L1 norm between the KOMB profiles of the subjects. Supplementary Figure S6 shows the pairwise distances as calculated by the proposed measure. To get a better estimate of the difference between each probability distribution we grouped samples from three of the subjects Alien, Bugkiller, and Peacemaker according to time points, namely initial comprising Days 0, 2, 7, and 60, post-antibiotic comprising Days 376, 377, 378, and 380, and only from Alien and later comprising Days 392 (3 samples) and 773. This grouping was motivated by the hypothesis that the distance between Alien initial and Alien post-antibiotic was significantly greater than a change that could be explained merely by a difference in time duration. From Supplementary Figure S6 we indeed observe that Alien post-antibiotic has significantly greater pairwise distance to all other samples (Avg dist = 0.622). This also happens to be far more than the distance between samples of subjects at initial and later time points (Avg dist = 0.312). Observing samples collected from Alien, the average pairwise distance between Alien initial and other samples (excluding Alien post-antibiotic) is 0.227 and that between Alien later and other samples (excluding Alien post-antibiotic) is 0.38. No statistical testing was done given the novelty of these metrics and small number of data point, but nonetheless the distances appear to reinforce the conclusion that antibiotic intervention does in fact cause significant perturbation in KOMB profiles.

### FMT samples pre, post, and donor

We analyzed two patient samples at two different time-points namely, Pre-FMT and Post-FMT using KOMB to understand shift in microbiome communities after an FMT procedure. We also compared the KOMB anomaly profiles of Pre-FMT and Post-FMT samples to the Donor sample to track common patterns between them. The Pre-FMT samples were collected from the patients post vancomycin treatment. In Figure 6(A), we observe that the anomaly profiles of Pre-FMT samples are distinctly shrunk (less coreness) compared to the Post-FMT and Donor samples indicating similar trends previously observed after the antibiotic treatment in the gut microbiome study [51]. We also see that Patient 1 shows some partial recovery towards the Donor profile whereas Patient 2 shows a higher similarity to the Donor in terms of coreness and anomaly score.

**Figure 6.**
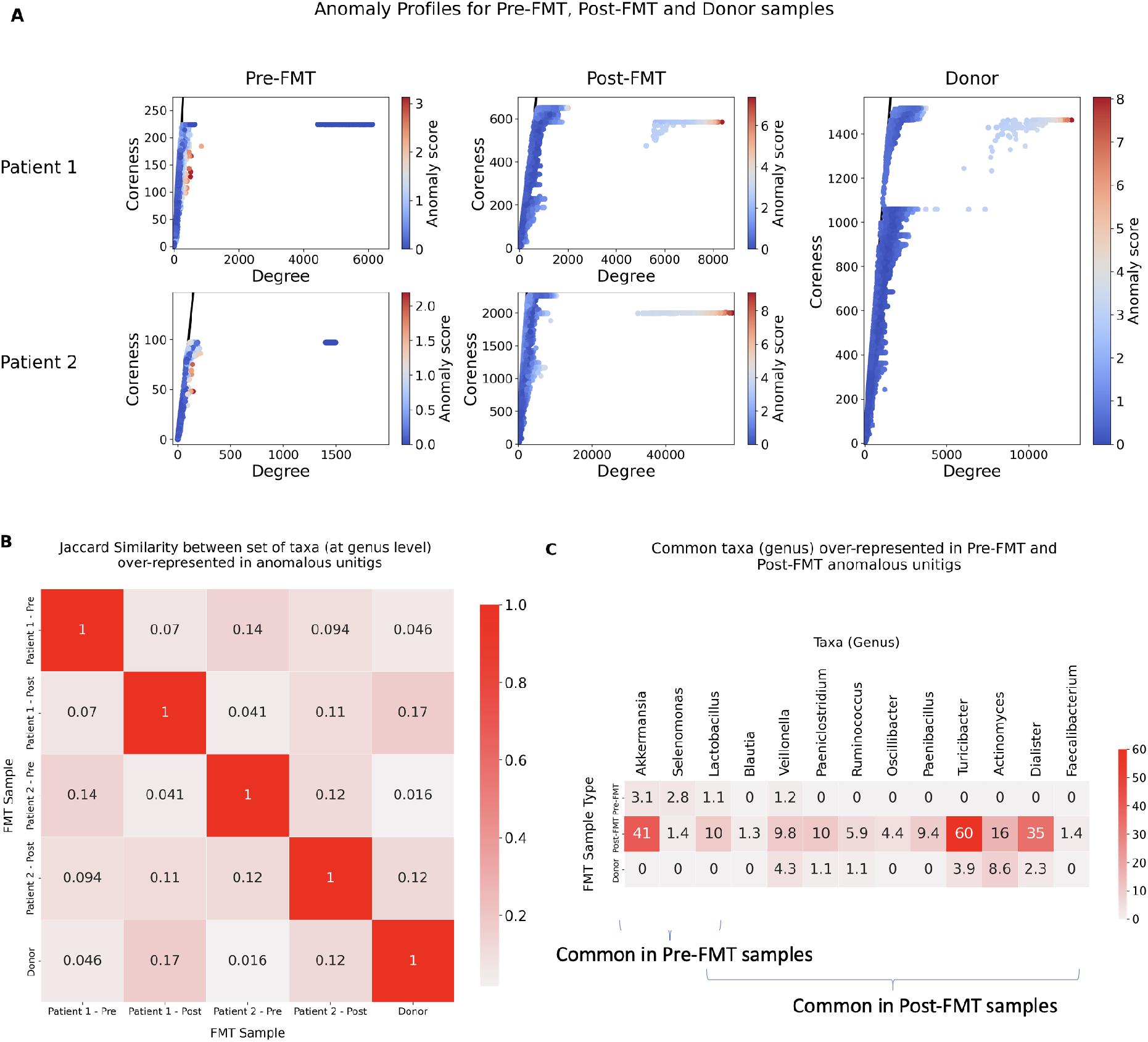
Characterizing community shifts in fecal microbiota transplantation (FMT) samples. (A) (Left) Anomaly profiles o f t wo p atients u ndergoing F MT t herapy a t t wo different time points namely Pre-FMT and Post-FMT. (Right) Anomaly profiles of the donor sample, which is common for both patients. The x-axis represents the degree of unitigs and y-axis represents the coreness (or shell number) of the unitigs. The gradient on the colorbar indicates the CORE-A anomaly scores of unitigs in the sample. (B) Jaccard similarity between sets of taxa over-represented at the genus level found in unitigs marked as anomalous in each of the 5 samples. The row highlighted in black indicates the jaccard similarities of each patient across time points as compared to Donor. (C) Common taxa over-represented in anomalous unitigs for Pre-FMT, Post-FMT and Donor samples. The numbers indicate the ratio of ratios of counts of taxa, indicating the relative level of presence of the corresponding taxa in the anomalous unitig compared to the other unitigs in the sample. The numbers in the figures have b een averaged for Pre-FMT and Post-FMT samples from both Patients. The first three genus *Akkermansia*, *Selenomonas* and *Lactobacillus* were common in Pre-FMT while *Lactobacillus*, *Blautia*, *Veillonella*, *Paeniclostridium*, *Ruminoccocus*, *Oscillibacter*, *Paenibacillus*, *Turicibacter*, *Actinomyces*, *Dialister*, *Faecalibacterium* were common in Post-FMT samples.

The unitigs obtained after KOMB analysis from one of the Post-FMT samples were too short and fragmented to annotate functionally using SeqScreen. In lieu of this, we examined the taxa represented by anomalous unitigs with the thinking that they may indicate important organisms driving the change in host microbiome post-FMT. For unitigs identified as anomalous (and which could be classified at the genus level, See Methods), over-represented taxa were determined by the score defined in Equation 4. In Figure 6(B) we see that, in general, there is a low similarity between over-represented taxa across the samples. We still observe that for both Patients 1 and 2, the Post-FMT samples have a higher taxa similarity to Donor compared to the Pre-FMT samples (highlighted by the black box in Figure 6(B)) as captured in the anomalous unitigs despite a substantial difference in their anomaly profiles.

As seen in Figure 6(C), Pre-FMT samples had three genera in common; *Akkermansia*, *Selenomonas* and *Lactobacillus* whereas Post-FMT had eleven: *Lactobacillus*, *Blautia*, *Veillonella*, *Paeniclostridium*, *Ruminoccocus*, *Oscillibacter*, *Paenibacillus*, *Turicibacter*, *Actinomyces*, *Dialister*, *Faecalibacterium* in common. We compared the relative levels of these taxa in Pre-FMT and Post-FMT and Donor. The values in the heatmap represent the average of the Ratio of Ratios score in both patients. Compared to Pre-FMT levels, we saw a substantial increase in two taxa *Akkermansia*and *Lactobacillus* in the Post-FMT anomalous unitigs. *Akkermansia* was only overexpressed in Patient 2, while *Lactobacillus* was found in both Patient 1 and Patient 2 at Post-FMT. Further analysis showed that *Akkermansia* was also present in anomalous unitigs in Patient 1 but at a much lower level than the background. Interestingly, previous studies have shown that higher levels of some species belonging to *Akkermansia* and *Lactobacillus* were helpful to combat *Clostridium difficile* infections [61, 62]. In contrast to Pre-FMT and Post-FMT *Akkermansia*, *Selenomonas* and *Lactobacillus* were also present in the anomalous unitigs in the Donor sample but were not over-represented compared to the other (background) unitigs.

Among the taxa common in Post-FMT Samples, roughly half (6/11) were similarly over-represented in Donor sample anomalous unitigs, though the levels were much higher in the former. However, *Turicibacter* and *Dialister* had a highest level of over-representation. This is noteworthy because *Turicibacter* is a well characterized taxa which is one of the most abundant in other reported studies on FMT inoculums and Post-FMT communities [63, 64, 65] whereas the presence of *Dialister* has been found to be essential in Post-FMT recovery and non-disease states [66, 67]. Kraken 2 outputs and unitig classifications can be found in **Supplementary Data SD4**.

### Performance

KOMB is written in C++. It uses the igraph C graph library [41] for the unitig construction and K-core decomposition implementations. Table 1 shows the runtime and memory usage of KOMB on the datasets used in our study. The experiments were run on a server with 64 Intel(R) Xeon(R) Gold 5218 CPU @ 2.30GHz processors having 372 GB of RAM. While analyzing the runtimes of specific stages of the KOMB pipeline we observed that the ABySS unitig generation is the most memory intensive step in the pipeline while read mapping using Bowtie2 is the most computationally intensive step in the pipeline. As KOMB is also extremely memory efficient, one can process multiple metagenomic samples simultaneously on any modern workstation to reduce the runtime on entire datasets even further.

**Table 1.** Time and memory usage for KOMB. Shakya: Shakya et al (2013); HMP (Av); average across HMP samples, TGM(Av); average across Temporal Gut Microbiome samples and FMT (Av); average across FMT samples. Read filtering is treated as a pre-processing step, therefore the time and memory usage for it is not reported in this table. KOMB was run with 20 threads.

| Dataset | Performance metrics |  |  |  |  |  |
| --- | --- | --- | --- | --- | --- | --- |
|  | *Reads* | *Nodes* | *Edges* | *Wall clock* | *CPU time* | *RAM* |
| Shakya | 53,997,046 | 96,901 | 1,080,012 | 77m50s | 1296m21s | 25.29 GB |
| HMP (Av) | 16,872,599 | 303,171 | 2,414,541 | 26m34s | 445m1s | 9.59 GB |
| TGM (Av) | 26,520,076 | 776,058 | 7,286,158 | 44m41s | 810m48s | 20.22 GB |
| FMT (Av) | 34,173,634 | 323,431 | 22,994,009 | 93m56s | 1576m38s | 23.16 GB |

## Discussion

We have underlined the usefulness of characterizing metagenomes with KOMB through three separate use-cases. First, KOMB can be used to obtain a repeat profile of a sample capturing the more repetitive unitigs in the later shells. Second, KOMB can be used to identify and extract functionally rich unitigs and visualize sample-specific or subject-specific profiles as observed in our analysis of HMP samples from four distinct body sites. Finally, we also show how KOMB profiles can help with analyzing community shifts and disruption events in longitudinal samples.

Though KOMB is not intended to be a repeat detection tool for metagenomes, it offers a convenient way of obtaining a sample-wide profile of repetitive unitigs. Repeat detection in metagenomes is a complex task and several previous methods have attempted to identify repeats and/or repeat families through different methods. KOMB attempts to identify sequences containing a small subset of exact repeats of length equal to k-1 where k is the k-mer size used to build the De Bruijn graph. One advantage of KOMB is that, given our hybrid unitig graph construction, the K-core decomposition gives a sample wide profile of exact repeats depending on their copy number. The hybrid unitig graphs draws inspiration from other previously studied De Bruijn graph types with embeddings or support for efficient repeat retrieval like A-bruijn graphs [68],Linked De Bruijn Graphs [69] and SIGAR graphs [70]. As seen by our results in the synthetic metagenome dataset, denser repeats are likely to be found in the higher shells. These could refer to certain inter-genomic repeats or intra-genomic repeats based on their relative copy numbers as well as the number of genomes the unitig is shared by. In addition to this, a single shell could contain multiple unitigs representing different repeat families if they share the same copy number (or occur in similar parts of the genome). A natural extension of this work would be to incorporate inexact repeats to the existing framework, which would help capture more biologically relevant relationships and account for subtle homology differences between regions of similar organisms at the cost of a longer runtime.

KOMB leverages the underlying topology of the hybrid unitig graph to use the K-core graph decomposition algorithm to identify anomalous unitigs that could identify functionally rich unitigs. These anomalous unitigs had the highest CORE-A score. Nodes having high CORE-A score exhibit high coreness and low degree or low coreness and high degree. The latter resembles topologies picked up by betweenness centrality measures described in previous works [28, 29, 30]. Identification of unitigs having high coreness and low degree is a unique feature of KOMB that enables identification of regions of dense repeats in a metagenome. A significant advantage KOMB offers in comparison to centrality based methods is the favourable *O*(*E + V*) runtime to identify both kinds of anomalous unitigs. Another advantage is that KOMB performs a de-novo decomposition of the hybrid unitig graph and does not depend on user-defined neigborhood queries or taxonomic labels to characterize samples. Results on the HMP data from four body sites show that functional enrichment of the anomalous unitigs highlight important functional differences between the communities in each sample. KOMB was able to capture unique functional terms at a statistically significant level which could be useful to generate functional summaries of microbiome communities.

Through taxonomic validation on our analysis of the KOMB profiles of FMT samples, we were able to show how the unitigs marked as anomalous can potentially belong to species indicative of the condition of FMT patients. Genera over-represented in anomalous unitigs in the Post-FMT samples were indicative of a transitional shift in the microbiome community as compared to Pre-FMT (Vancomycin treated) and Donor Samples underlying KOMB’s usefulness in summarizing community shifts in loongitudinal samples.

KOMB, to the best of our knowledge, represents the first method to unify the extraction of graph based topological features and k-mer based methods to characterize metagenomes. Compared to previous graph based methods, KOMB offers the ability to visualize and calculate intra-sample distances. Compared to k-mer based methods, KOMB allows for de-novo analysis and extraction of functionally rich as well as relevant taxonomic sequences in metagenomic samples. Despite its strengths, there are some natural future enhancements that could be explored. First, KOMB is slower and much more memory intensive than some of the k-mer based methods. While some of this cost is necessary to gain a more sequence level view of the sample, other efficient (or lossy) De Bruijn graph constructors could be considered to make the process more scalable to extremely large metagenomic samples with billions of reads. Second, KOMB relies exclusively on topology suited for retrieval by K-core decomposition i.e it relies on extracting clique or clique-like regions that are connected due to repeats or paired-end information. Future work to analyze other biologically relevant topologies that can be extracted by hybrid unitig graphs or its variants of the graph could be useful. Third, K-core profile of the unitigs can be used to obtain an approximate value of the entropy in samples, KOMB still lacks a direct conversion to popular diversity measures as provided by other k-mer based approach which would need further theoretical analysis.

## Conclusions

In summary, KOMB can be used to obtain sample-wide repeat profiles, visualize community shifts and disruption events in longitudinal gut microbiome samples, and quantify inter-sample distances across various time points. Combined with its ability to identify sample-specific and biologically important unitigs, KOMB can be used to get a holistic characterization of metagenomic samples both at a macro(sample) as well as at micro(sequence) level.

## Declarations

### Ethics approval and consent to participate

The FMT samples were obtained under IRB-approved informed consent (#H-31066) at Baylor College of Medicine.

### Competing interests

The authors declare they have no competing interests.

### Availability of data and materials

Python jupyter notebooks used for analysis and generating figures can be found here: https://rb.gy/zxrv1z. KOMB outputs and files used for analysis for the experiments can be found at https://tinyurl.com/f42t96rb

### Availability and requirements

The latest version of KOMB (v1.0) is also available for download through bioconda at: https://anaconda.org/bioconda/komb

Project name: KOMB

Project home page: https://gitlab.com/treangenlab/komb

Operating system(s): OSx and Linux Programming language: C/C++

Other requirements: Requirements installed as part of bioconda install. At least 64GB of RAM recommended.

License: GNU GPL v3.0 or later

Any restrictions to use by non-academics: No

### Authors contributions

A.B, T.J.T, S.S developed the study. A.B implemented the software, performed the validation, analyzed the data, interpreted the results, generated figures and wrote the manuscript. N.S performed the validation, analysed the data and generated figures. C.S analyzed the data and interpreted the results. R.A.L.E, S.S, T.S and T.J.T contributed to the design of the validation and the interpretation of the results. M.G.N helped with the writing of the manuscript. All authors read and approved the final manuscript.

### Funding

A.B. and T.J.T were supported by startup funds from Rice University and the FunGCAT program from the Office of the Director of National Intelligence (ODNI), Intelligence Advanced Research Projects Activity (IARPA), via the Army Research Office (ARO) under Federal Award No. W911NF-17-2-0089. R.A.L.E. was supported by the FunGCAT program from the Office of the Director of National Intelligence (ODNI), Intelligence Advanced Research Projects Activity (IARPA), via the Army Research Office (ARO) under Federal Award No. W911NF-17-2-0089. N.S is supported by Department of Computer Science, Rice University. C.S and T.S were supported by the P01-AI152999 and U01-AI24290 grants obtained from the National Institutes of Health (NIH). M.N. was supported by a fellowship from the National Library of Medicine Training Program in Biomedical Informatics and Data Science (T15LM007093, PI: Kavraki).

The views and conclusions contained herein are those of the authors and should not be interpreted as necessarily representing the official policies or endorsements, either expressed or implied, of the ODNI, IARPA, ARO, or the US Government.

## Acknowledgments

The authors would like to thank Dr. Mihai Pop and the Pop Lab at University of Maryland, College Park for their constructive comments and suggestions that helped refine the manuscript.

## Notes

https://gitlab.com/treangenlab/komb

### Competing Interest Statement

The authors have declared no competing interest.

