## SUPPLEMENTARY DATA for "KOMB: Graph-Based Characterization of Genome Dynamics in Microbial Communities"

### Supplementary
### KOMB: Reference-free Characterization of Metagenome Dynamics via K-core Decomposition

Advait Balaji[1], Nicolae Sapoval[1], Charlie Seto[2], R.A. Leo Elworth[1], Michael G. Nute[1], Tor Savidge[2], Santiago Segarra[3] & Todd J. Treangen[1#]

[1] *Department of Computer Science, Rice University, Houston, Texas, USA.*
[2] *Department of Pathology and Immunology, Baylor College of Medicine, Houston, TX, USA*
[3] *Department of Electrical and Computer Engineering, Rice University, Houston, Texas, USA.*

#### Supplementary Data

**SD1.** L1 norm analysis of KOMB Profiles from HMP sites

We calculate the L1 norm of KOMB profiles both within and between each body site. For samples within in each body site, anterior nares had the highest average distance (1.15) followed by buccal mucosa (1.14), supragingival plaque (1.00) and stool (0.89). The aggregate distances (see implementation) between body sites were also calculated. We observed that anterior nares and buccal mucosa had the highest aggregate distance (1.09). Overall, anterior nares also had the greatest separation from supragingival plaque (1.08) and stool (0.66). Supragingival plaque and buccal mucosa were closest in terms of aggregate distance (0.32).

**SD2.** GO terms obtained from the HMP analyses can be found at: https://tinyurl.com/3s5fe99k

**SD3.** L1 norm analysis of KOMB Profiles fro Subjects in Gut-microbiome analysis

We calculated the average pairwise L1 norm for samples from each subject. Bugkiller (0.30), Scavenger (0.46), Tigress (0.34), and Daisy (0.38) showed higher variability in the early samples as compared to other subjects Alien (0.12) and Peacemaker (0.16) who exhibited fairly consistent profiles. We generally observe intra-sample similarity over the three time points and also observe some similarities between profiles based on gender also reported by previous studies [1, 2]. Aggregate profiles from Daisy and Tigress were closest to each other (0.26) than to any of the male subjects. The average distances of the male subjects to Daisy and Tigress were; Alien (0.58, 0.33), Bugkiller (0.62,0.37), Peacemaker (0.61,0.36) and Scavenger (0.52,0.27) respectively while the average pairwise distance between profiles from the male subjects was 0.12.

**SD4.** Kraken2 analysis on FMT samples can be found at: https://tinyurl.com/yhr9w8hv

### References

[1] JA Santos-Marcos, C Haro, A Vega-Rojas, JF Alcala-Diaz, H Molina-Abril, A Leon-Acuña, J Lopez-Moreno, BB Landa, M Tena-Sempere, P Perez-Martinez, et al. Sex differences in the gut microbiota as potential determinants of gender predisposition to disease. *Molecular nutrition & food research.* **63**: 1800870.

[2] F Fransen, AA van Beek, T Borghuis, B Meijer, F Hugenholtz, C van der Gaast-de Jongh, HF Savelkoul, MI de Jonge, MM Faas, MV Boekschoten, et al. The impact of gut microbiota on gender-specific differences in immunity. *Frontiers in immunology.* **8**: 754.

### Supplementary Figures

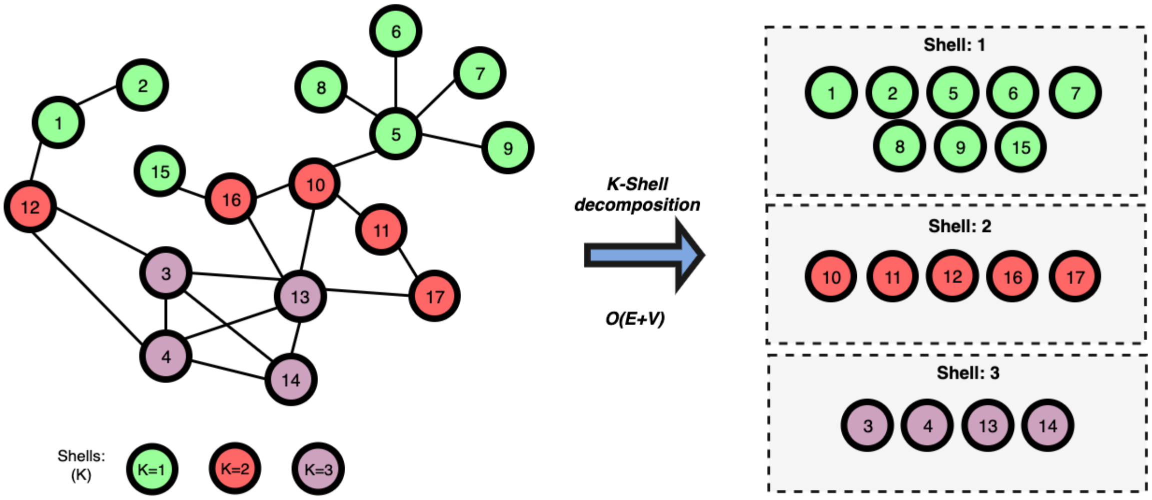


**Figure S1. K-core decomposition of a graph into K-shells.** The algorithm starts by considering all the vertices of degree 1. It iteratively removes those vertices and continues the execution on the resulting induced subgraph removing vertices having degree 1 after every iteration. Once the induced subgraph has no vertices of degree 1, this process stops and all discarded vertices are marked as belonging to the 1-shell (green). Then the process continues, now considering vertices of degree 2 to obtain the 2-shell (red) and, subsequently, the 3-shell (purple). The last shell is a dense subgraph of the original graph.

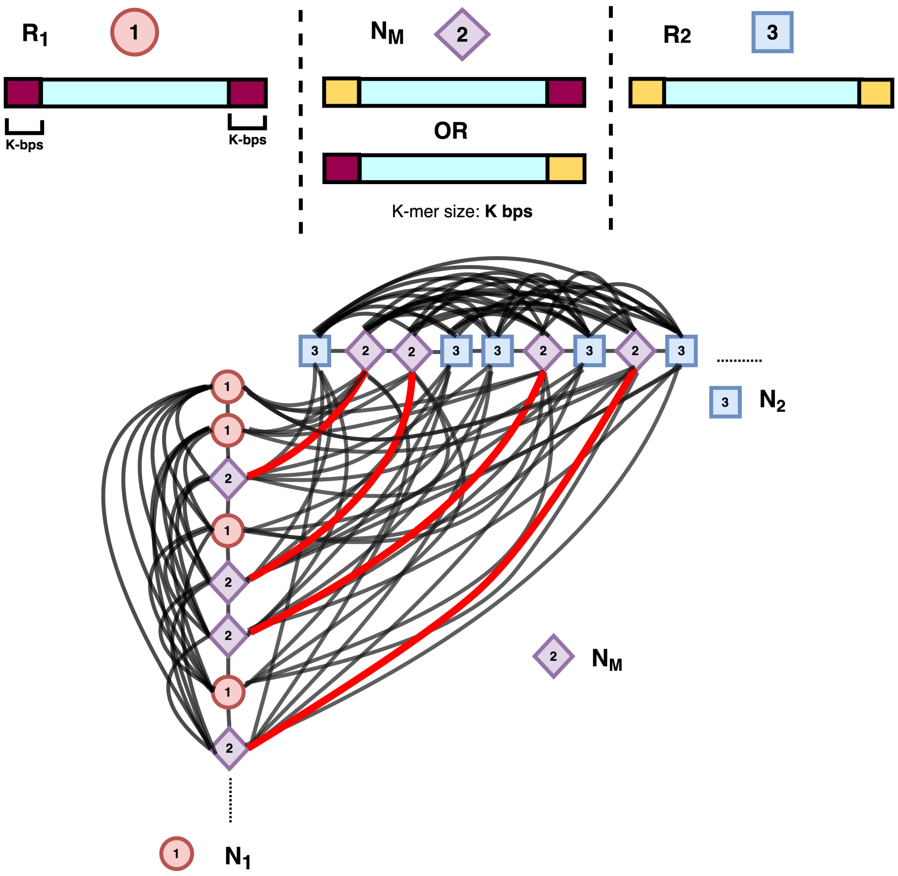


**Figure S2. Types of unitigs in a genome with two repeat families and expected shell profiles in corresponding unitig graphs.** The type of profile we observe depends on the relative lengths of the repeats and insert size. If the insert size is greater than the length of the repeat, the mixed repeats ($N_m$) will be connected to each other whereas if the insert size is smaller than the length of the repeat then it is not possible to map across the two mixed repeat unitigs and, hence, they will not be connected by an edge in the unitig graph. The black edges are present for both settings whereas the red edges are only present when the repeat length is less than the insert length.

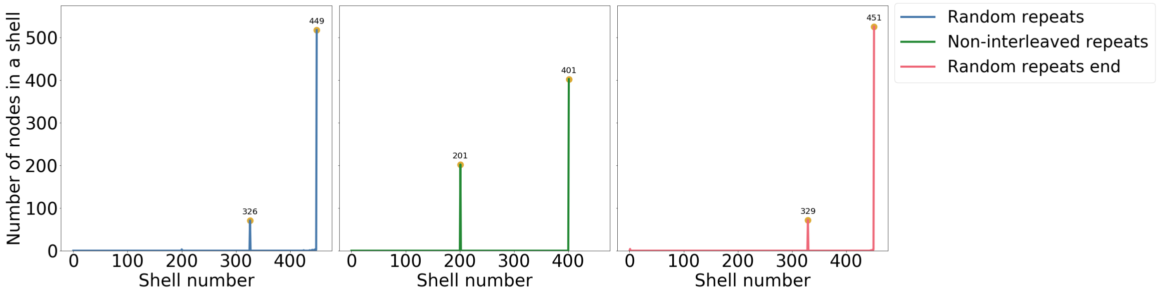


**Figure S3. Validation of KOMB on simulated data.** KOMB profiles on a random backbone with 200×400bp and 400×200bp identical repeats when randomly inserted (L), inserted at different ends (M), and inserted in the same end (R). The x-axis represents the shell number and the y-axis represents number of nodes (unitigs). As discussed in Methods, interleaving of repeats, as observed in (L) and (R) creates unitigs with different repeats at it ends. In the ideal case (M) where the unitigs have the same repeats at its end we observe peaks exactly at 200 and 400 respectively. Interleaving repeats randomly (L) causes a shift in the peaks towards higher shells to 326 and 449 respectively. Further, inserting both kinds of repeats at the same end (R) results in an even higher shift with peaks occurring at 329 and 451.

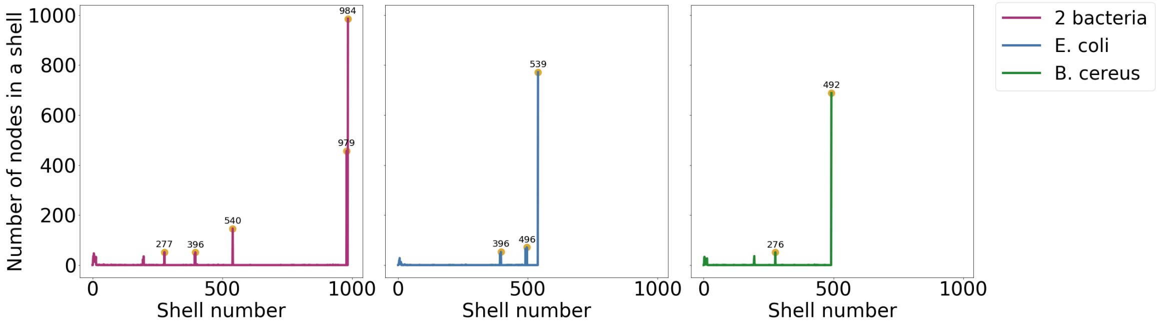


**Figure S4. Validation of KOMB on simulated data with a real genomic backbone.** Combined KOMB profile of *E. coli* (intra: 400×400bp, inter 500××500bp) and *B. cereus* (intra: 200×400bp, inter 500×500bp) repeats (Left), *E. coli* single genome (Middle), and *B. cereus* single genome (Right). For the combined profile of both bacteria (L), there is a clear formation of peaks close to position 1000 (984 and 979), which indicate the inter-genomic repeats, and peaks at shell numbers 277, 396 and 540. This agrees with the theoretical model in the case of unitigs with mixed repeats at its end as discussed in Methods. We also plot the individual profiles of *E.coli* and *B.cereus*. For *E.coli* (M) we see three peaks at 396, 496 and, 539 given its higher copy number of intra-genomic repeats (400). For *B.cereus*, we observe peaks at 276 and 492 signalling its intra and inter-genomic repeats, respectively.

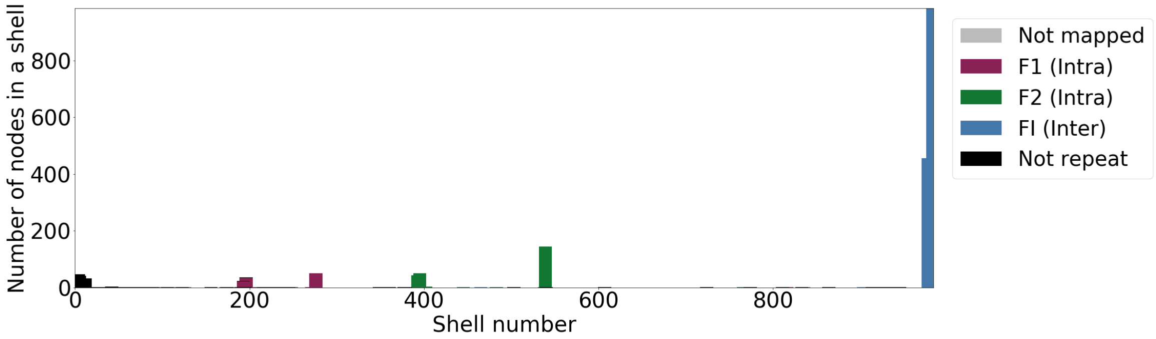


**Figure S5. Validation of repeat types in *E. coli* + *B. cereus* sample via mapping unitigs back to the reference using nucmer.** Unitigs are labelled based on the repeats they overlap with. Based on ground truth nucmer mapping, the last shell (close to 1000) contains unitigs overlapping exclusively with inter-genomic repeats (FI) whereas the shells around 200 are overlapping *B. cereus* simulated intra-genomic repeats (F1) and shells around 400 are overlapping *E. coli* simulated intra-genomic repeats (F2). Finally, the first shells contain the background noise (colored black).

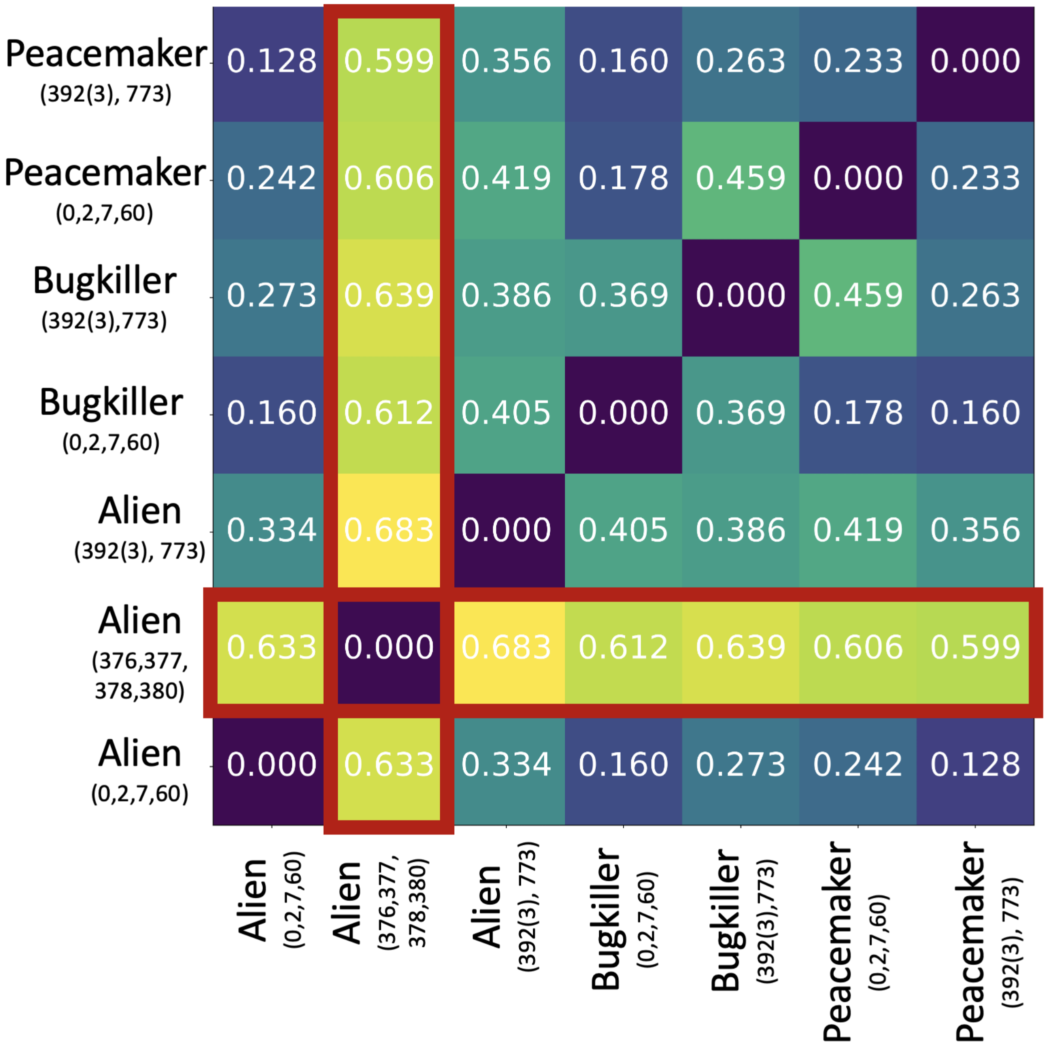


**Figure S6. Heatmap showing L1 norm of KOMB profiles in the Voigt et al. dataset.** Each row and column represents a subject and days (four each) for which the samples are considered (in parenthesis). Day 392 had 3 samples in the dataset which are all considered here. The samples represented by Alien (Days 376, 377, 378, and, 380), also marked in red, are the ones collected during antibiotic perturbation. Higher total variation of probability denotes greater distance between two distributions. Days 0,2,7,60 correspond to the initial time points and Days 392(3) and Day 773 correspond to later time points.
